# Analysis of human immunoglobulin VDJ and DJ rearrangements shows N region synthesis by concatenation of cytosine-rich strands preferentially originating from trimmed germline gene segments

**DOI:** 10.1101/248021

**Authors:** Tina Funck, Mike Bogetofte Barnkob, Nanna Holm, Line Ohm-Laursen, Camilla Slot Mehlum, Sören Möller, Torben Barington

## Abstract

The formation of non-templated (N) regions during immunoglobulin gene rearrangement is a major contributor to antibody diversity. To gain insights into the mechanisms behind this, we studied the nucleotide composition of N regions within 29,962 unique human V_H_DJ_H_-rearrangements and 8,728 unique human DJ_H_-rearrangements containing exactly one identifiable D-gene segment and thus two N regions, N1 and N2. We found a distinct decreasing content of cytosine (C) and increasing content of guanine (G) across each N region, suggesting that N regions are typically generated by concatenation of two 3’-overhangs synthesized by addition of nucleoside triphosphates with a preference for dCTP. This challenges the general assumption that the terminal deoxynucleotidyl transferase favors dGTP *in vivo*. Furthermore, we found that the G and C gradients depended strongly on whether the germline gene segments were trimmed or not. Our data show that C-enriched N addition preferentially happens at trimmed 3’-ends of V_H_-, D-, and J_H_-gene segments indicating a dependency of the transferase mechanism upon the nuclease mechanism.

## Introduction

Diversity in the immunoglobulin repertoire is paramount for the immune system’s ability to fight the multitude of foreign pathogens that the body encounters. Diversity if created through random combination of one of many V, (D), and J gene segments giving rise to the V(D)J genes encoding the variable domains of B- and T-cell receptors. However, the major contributor to diversity is the variable truncation of the recombined gene segments in synergy with the addition of non-templated nucleotides forming the so-called N regions within the rearrangements (1). In human B cells, N regions are usually present in both heavy and light chain genes in fetal as well as adult B cells, whereas they are only present in heavy chains in murine B cells and only after birth (2–4). These discrepancies are caused by different timing of the expression of Terminal Deoxynucleotidyl Transferase (TdT), an enzyme directly involved in the formation of N regions (5, 6). Initiating the recombination process, Recombination-activating gene 1 (RAG1) and RAG2 induce a double-stranded DNA break between the gene segment and the flanking recombination signaling sequence (RSS) and seal the break with a hairpin loop (7, 8). Initially, this happens for a random D and J_H_ gene segment pair on one of the human chromosome 14s. Later the same happens between a random V_H_ gene and the formed DJ_H_ gene. In both cases, Ku70/Ku80 heterodimers bind to the DNA ends and recruit the Artemis:DNA-PKcs complex to which also XRCC4 associates (9). The endonuclease Artemis then nicks the DNA strand at variable distance from the sealed end, which usually results in formation of 3’-overhangs at the two gene segments (10). These overhangs sometimes remain in the final rearrangement as an extension of the gene segment(s) known as palindromic nucleotides (P segment), but often nucleotides are removed leading to trimming of the germline encoded gene segment(s). Unique for the recombination events occurring on the B- and T-cell receptor loci is the involvement of TdT (11–13). TdT is only expressed in the early developmental stages of B and T cells. TdT works on the exposed 3’-overhangs and has both exonuclease activity and nucleotidyl transferase activity. One short and two long splice variants of TdT have been identified in humans and *in vitro* studies using purified human TdT have indicated that the short variant executes the transferase activity while the two long variants have 3’-5’ exonuclease activity (14, 15). The exonuclease activity of TdT is only effective on 3’-overhangs and does not progress into double stranded DNA (16). Thus both TdT and Artemis could be involved in the elimination of the single-stranded 3’-ends, TdT by its single-strand exonuclease activity and Artemis by its endonuclease activity. TdT may extend 3’-overhangs as well as 3’-ends of blunt-end DNA. Eventually, the free single-stranded 3’-overhangs on the two gene segments to be joined will pair guided by sequence micro-homology by the process non-homologous end joining (NHEJ) which involves the nuclease APLF, the DNA ligase IV complex, and the DNA polymerases λ and µ (17, 18), for review see: (19). Mismatching nucleotides are removed and gaps are filled by DNA polymerases λ and µ (20–22). While it is known that TdT interacts with the DNA-PKcs complex (23), it is unclear whether TdT adds nucleotides on the plus strand, the minus strand or both. Kepler *et al.* and others (24, 25) have suggested that murine N regions are predominantly formed by a single 3’-overhang growing from the plus strand (here designated the single-strand polymerization hypothesis), whereas in humans it has long been proposed that N regions are derived by concatenation of two 3’-overhangs (here designated the concatenation hypothesis) (11). Both hypotheses were proposed based on analyses of a limited number of rearrangements and neither has been firmly established. TdT has preference for certain nucleotides, and hence a strand bias for TdT activity will heavily affect the resulting antibody repertoire. It remains unexplored whether the N region between the V_H_- and D-gene segments (N1) and the D- and J_H_-gene segments (N2) are constructed similarly and whether TdT adds nucleotides to the upstream, downstream or at both recombining gene segments.

In this paper, we explore the nucleotide composition of N regions in a large dataset consisting of 29,392 productive (P) and 4,708 non-productive (NP) V_H_DJ_H_-rearrangements and 9,061 sterile DJ_H_-rearrangements, all with a single D-gene segment. The dataset thus includes 13,769 rearrangements from which we have been able to study the nucleotide composition with no bias from antigen selection. Furthermore, 12,499 of these rearrangements were unmutated and therefore likely to represent the preferences of the rearrangement machinery *per se*. From this dataset, we are able to infer several conclusions regarding the strand usage, nucleotide preferences *in vivo* during N addition and the role of gene segment trimming.

## Materials and Methods

### Datasets

#### Sanger sequences

Two sets of unique V_H_DJ_H_-rearrangements were produced by PCR amplification of genomic, rearranged DNA from B cells using IGHV3-23-specific and IGHJ4 and −6 favoring primers followed by cloning and Sanger sequencing as previously described (26). Unique rearrangements were defined as sequences which differed from all other sequences in the material by more than four base positions in the joint region defined as bases aligned to the last 20 nucleotides of the germline V_H_-gene segment or to the J_H_ gene segment and all bases between. The first set of 1,886 unique V_H_DJ_H_-rearrangements was obtained from CD27 negative B-cell fractions from blood obtained from 10 healthy adults enrolled in a previous study (26). The other set comprised 1,695 unique V_H_DJ_H_-rearrangements obtained from palatine tonsils removed from 8 patients with chronic tonsillitis (3-42 years of age, 3 males). Individuals were enrolled after informed consent and the scientific ethics committee of Vejle and Funen Counties approved the studies (S-VF-20040071 and VF-20030161).

The tonsils were mechanically homogenized in isotonic saline to prevent cell lysis. After sedimentation of debris, mononuclear cells were isolated using density gradient centrifugation over Lymphoprep (Axis-Shield, Roskilde, Denmark) and DNA was extracted from the PBMCs.

#### Next generation sequences

A third dataset of 16,893 unique V_H_DJ_H_-rearrangements was obtained by next generation sequencing from B cells purified from anonymized buffy coats prepared from blood donations by 10 healthy blood donors in agreement with Danish legislation. The donors gave written consent for their blood to be used for research purposes. PBMCs were purified using lymphoprep (Axis-Shield) and equal numbers of PBMCs from 5 donors were pooled forming two pools of PBMCs. B cells were purified from both by negative selection using EasySep™ Human B Cell Enrichment Kit (Stemcell Technologies) from 450*10^6^ PBMCs. The B cells were subsequently positively purified with regard to their expression of either kappa or lambda light chain. This was carried out using CELLection (Life Technologies) magnetic beads pre-coated with murine monoclonal IgG specific for either human kappa or lambda light chains (BD Bioscience). DNA was extracted by Maxwell 16 Blood DNA purification kit (Promega) and the DNA concentrations measured by Qubit dsDNA BR Assay Kit (Invitrogen). V_H_DJ_H_-rearrangements were PCR amplified using one of 9 primers covering the 7 V_H_-gene families (forward) and a mixture of 3 reverse primers covering the 6 J_H_-genes. The following forward primers were used: IGHV1: 5’-TGAGGTGAAGAAGCCTGGG-3’, IGHV2 5’-CTCTGGGTTCTCACTCAGC-3’, IGHV3-1: 5’-GTCCCTGAGACTCTCCTGT-3’, IGHV3-2 5’-GAGGTGCAGCTGGTGGAG-3’, IGHV4 5’-ACCCTGTCCCTCACCTGC-3’, IGHV5: 5’-TGGAGCAGAGGTGAAAAAGC-3’, IGHV6: 5’-TTGCTGTTTCCTTTTGTCTCC-3’, IGHV7(1): 5’-GCTTCTGGATACACCTTCAC-3’, IGHV7(2): 5’-ATGAATTGGGTGCGACAGG-3’ and reverse primers: IGHJ1,4,5,6: 5’-CCTGAGGAGACGGTGACC-3’, IGHJ2: 5’-CCTGAGGAGACAGTGACCA.3’, IGHJ3: 5’-CCTGAAGAGACGGTGACCA-3’. The PCR was run using AmpliTaq Gold (Life Technologies) in a total volume of 25 µL using 50 ng of genomic DNA, 2.5 mM dNTP, 0.250 µM forward primer, and 0.125 µM of each of the IGHJ2 and IGHJ3 primers and 0.25 µM of the IGHJ1,4,5,6 primer. The PCR was run with initial denaturation at 96⁰C for 12 minutes followed by 35 cycles of denaturation at 96⁰C for 30 seconds, annealing at 60⁰C for 30 seconds, and elongation at 72⁰C for 30 seconds, followed by a final elongation step for 7 minutes. The amplicons were checked by gel electrophoresis. Library preparation was carried out using the Fragment Library Kit (Life Technologies) as instructed by the manufacturer and using AMPure beads for library clean up (Beckman-Coulter). Concentration measurements were done by Qubit dsDNA HS Assay Kit (Invitrogen), in some cases supplemented by Bioanalyzer Agilent High Sensitivity DNA Kit (Agilent Technologies). Libraries derived from the same pool of five donors and the same light chain were pooled after adjustment to the same amount of PCR product (2 nM) and underwent end-repair, ligation of barcoded adapters (Ion Xpress Barcode Adapters) and amplification as recommended by the manufacturer (Life Technologies). Separate MIDs were chosen for the four libraries. Isothermal amplification was performed using the Ion PGM Template IA 500 Kit according to the manufacturer’s protocol. Next Generation Sequencing was done on the Ion Torrent PGM platform using Ion PGM Hi-Q View sequencing Kit (Life Technologies). DNA from each pool of 5 donors was sequenced per Ion 314 chip v2. A total of 856,872 reads were analyzed using in-house software as previously described (27).

#### Published datasets

We also included two datasets previously published from our institution with broad representation of IGHV and IGHJ alleles. These comprised 5,800 unique V_H_DJ_H_-rearrangements and 9,969 unique sterile DJ_H_-rearrangements obtained by 454-sequencing and derived from CD19^+^ B cells isolated from 110 healthy blood donors (27).

Finally, we downloaded 72,107 putative V_H_DJ_H_-rearrangements from the NCBI repository (25^th^ of January, 2015) using IgBLAST and excluded entries containing the words: musculus, mouse, kappa light and lambda light, or those with ambiguous nucleotides in the sequence. Of these, 63,462 V_H_DJ_H_-rearrangements were classified as evaluable, meaning they had identifiable V_H_- and J_H_-genes and 39,992 of the rearrangements were unique with 32,064 being available in XML-format and linked to an NCBI Pubmed article. Based on data mining, we excluded rearrangements that originated from phage display, hybridomas, cell lines, mouse-human fusions, retracted papers *etc*. and ended up with an NCBI dataset including 24,411 unique V_H_DJ_H_-rearrangements.

In total, we gathered 50,685 V_H_DJ_H_ and 9,969 DJ_H_ sequences.

### V_H_DJ_H_-rearrangement analysis

We analyzed the V_H_DJ_H_- and DJ_H_-rearrangements in the program VDJsolver version 2.0. VDJsolver aligns rearrangements to the germline sequences of V_H_-, D-, and J_H_-genes using a maximum likelihood method (26). Version 1.0 is publicly available (http://www.cbs.dtu.dk/services/VDJsolver/) and has been described and benchmarked against other publicly available tools (28). Version 2.0 is described elsewhere (27) and differs from version 1.0 by: i) allowing D-D fusions in the model, ii) improved alignment with V_H_ germline genes using a Smith-Waterman-based algorithm (29) and iii) improved sensitivity for D-genes. The program found D-genes in 71% of the NCBI material and in 4% of artificially generated sequences in which empiric N(P)D(P)N stretches were replaced by stochastically assembled sequences with a base composition similar to N regions.

To assure precise definition of the N regions and to avoid short D segments to be misclassified as N nucleotides, sequences were only analyzed further if exactly one D-gene segment was found by VDJsolver.

VDJsolver splits V_H_DJ_H_-rearrangements using the model:

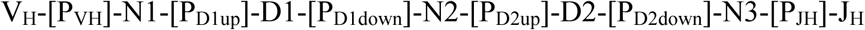

In the model “P” stands for palindromic segments at least two bases long immediately adjacent to an untrimmed gene segment end with suffixes designating the germline segment: V_H_, upstream or downstream end of the D segment, and J_H_, respectively. A single palindromic nucleotide immediately after an untrimmed gene segment was by default classified as an N nucleotide by VDJsolver, because the likelihood of an N nucleotide simulating a P nucleotide by chance exceeded the expected frequency of a single base P segment. To avoid any bias from P nucleotides misclassified as N nucleotides, such palindromic nucleotides were, however, ignored and not considered part of the N region in this paper. The need for precise definition of P segments required that the published (IMGT and PubMed) V_H_-gene germline sequence included the heptamer of the RSS to assure correct determination of the site of RAG-induced hairpin formation following double strand breakage. V_H_DJ_H_-rearrangements that used V_H_-genes with unresolved boundaries were therefore excluded from the study. These demands (V_H_ gene germline segment including heptamer and precisely one identifiable D gene in the rearrangement) reduced the sequence materials to 1,288 V_H_DJ_H_ rearrangements from tonsils (European molecular Biology Laboratory (EMBL) accession numbers XXX-XXX), 1,506 from CD27 negative B cells (accession numbers XXX-XXX), 4,083 V_H_DJ_H_- and 9,061 DJ_H_-rearrangements obtained by 454-sequencing (27), 11,381 V_H_DJ_H_ rearrangement obtained by Ion Torrent sequencing (accession numbers XXX-XXX), and finally 15,842 publically available V_H_DJ_H_ sequences; in total 43,161 informative sequences out of which 34,100 were V_H_DJ_H_- and 9,061 sterile DJ_H_-rearrangements. Nucleotide composition always refers to the plus strand.

### Identifying footprint sequences in N regions

Predicted N regions were screened for the existence of templated sequences compatible with secondary rearrangements – so-called footprints - which theoretically may be present in some of the N1 regions. All possible 7-mer, 6-mer and 5-mer motifs present between a putative cryptic RSS and the heptamer of the canonical RSS (Supplementary table I) in any of the known germline V_H_-genes were searched for in all N regions considering all possible starting positions as independent occurrences.

### Analysis of N regions

Except when explicitly mentioned, the first and the last nucleotide of each N region were ignored in the analysis of nucleotide composition of N regions, because these nucleotide positions were not free to attain any value (A, C, G, T) when classified by VDJsolver as N nucleotides. Some nucleotide values would have led to classification as part of the germline V_H_, D, J_H_ or a P segment. Nucleotide composition along N regions was studied for sequences with N regions ranging from 4-26 nucleotides in length and, ignoring the first and the last nucleotide, the individual nucleotide frequencies of the remaining 2-24 nucleotide positions were normalized by partitioning them into 10 deciles. Individual nucleotides were allowed to contribute to one or more deciles by the relevant fractions (Figure S1). For example, the fifth nucleotide of an N region of length 7 contributed by 0.2 in the 6^th^, 0.7 in the 7^th^, and 0.1 in the 8^th^ decile where the contribution to the 6^th^ decile was calculated as: (0.6−4/7)/ (5/7−4/7) = 0.2. Fractions were then summed up in each decile separately for all four nucleotides yielding a relative contribution of these nucleotides to each decile.

### Statistical analysis

Pearson’s chi-squared test was used when comparing numbers of the different nucleotides in N regions. Proportions of nucleotides as well as G/C-ratio across N regions were modelled by regression models weighted by number of observations and fitted by maximum likelihood estimation, and regression coefficients with 95% confidence intervals were estimated. The goodness of fit (*R*^2^) was estimated by weighted linear regression on the results of the maximum likelihood estimates. Slopes were interpreted as identical when regression coefficients (beta) did not deviate significantly from each other (*p*>0.05). The frequencies of putatively templated motifs in N regions were tested by Fisher’s exact test employing Holm-Bonferroni correction for multiple comparisons (30).

## Results

### N regions display linear gradients of G and C in the absence of selection

To study N regions as they were generated by the rearranging apparatus without the possible impact of antigen selection and somatic mutation, we restricted the first analysis to sterile DJ_H_-rearrangements and NP V_H_DJ_H_-rearrangements with unmutated V_H_-gene segments. In the following, the N region between the V-gene and D-gene segments is called N1 and that between the D-gene and J_H_-gene segments is called N2. Over all, the relative distributions of the four nucleotides, A, C, G, and T, did not differ between N1 and N2 prior to selection and mutation (A 20.9% vs. 20.7%; C 27.5% vs. 27.6%; G 31.2% vs. 30.7%; T 20.4% vs. 21.0%, *p*=0.226). Thus, G was the most common nucleotide and slightly but significantly more common than C in both N1 and N2 (*p*<0.0001). Heat maps revealed that G and C were not uniformly distributed in the N2 regions (Figure 1). Both nucleotides showed clear and opposite gradients from 5’- to 3’-end over a large range of N2 region lengths. In order to study these gradients more closely, the N regions ranging from 4-26 nucleotides in lengths were trimmed of their first and last nucleotide (for explanation, see M&M) and the remaining 2-24 nucleotides were divided into deciles according to their location within the N region. The fractions of A, C, G, and T nucleotides were determined for each decile as described in Materials and Methods and illustrated in Figure S1. For both N regions, clear linear gradients were found for the frequencies of G and C, while the frequencies of A and T were more uniformly distributed without significant slopes (Figure 2), except for a slightly negative slope for A in the N2 region (*R*^2^=0.52, *β*=−0.002 [-0.003;-0.001], *p*<0.0001). For N2, where the sample size was particularly large, the linearities of G (*R*^2^=0.97, *β*=0.009 [0.008;0.010]) and C (*R*^2^=0.96, *β*=−0.007[-0.008;-0.006]) frequencies were striking (*p*<0.0001). Even though the sample size was much smaller for N1, the linearity was convincing and significant (*p*<0.0001) for both the G (*R*^2^=0.80, *β*=0.006 [0.003;0.008]) and C (*R*^2^=0.92, *β*=−0.007 [-0.009;-0.004]). The opposite gradients for the G and C usage frequencies across N regions led to a considerable change in the G/C ratio throughout the N regions (Figure 2C). The G/C gradient observed in N1 (*R^2^*=0.95, *β*=0.048 [0.038;0.057], *p*<0.0001) and N2 (*R^2^*=0.96, *β*=0.063 [0.053;0.074], *p*<0.0001)) did, however, not differ significantly from each other (*p*=0.054). Neither did the A/T ratios (*p*=0.35) (data not shown).

**Figure 1.**
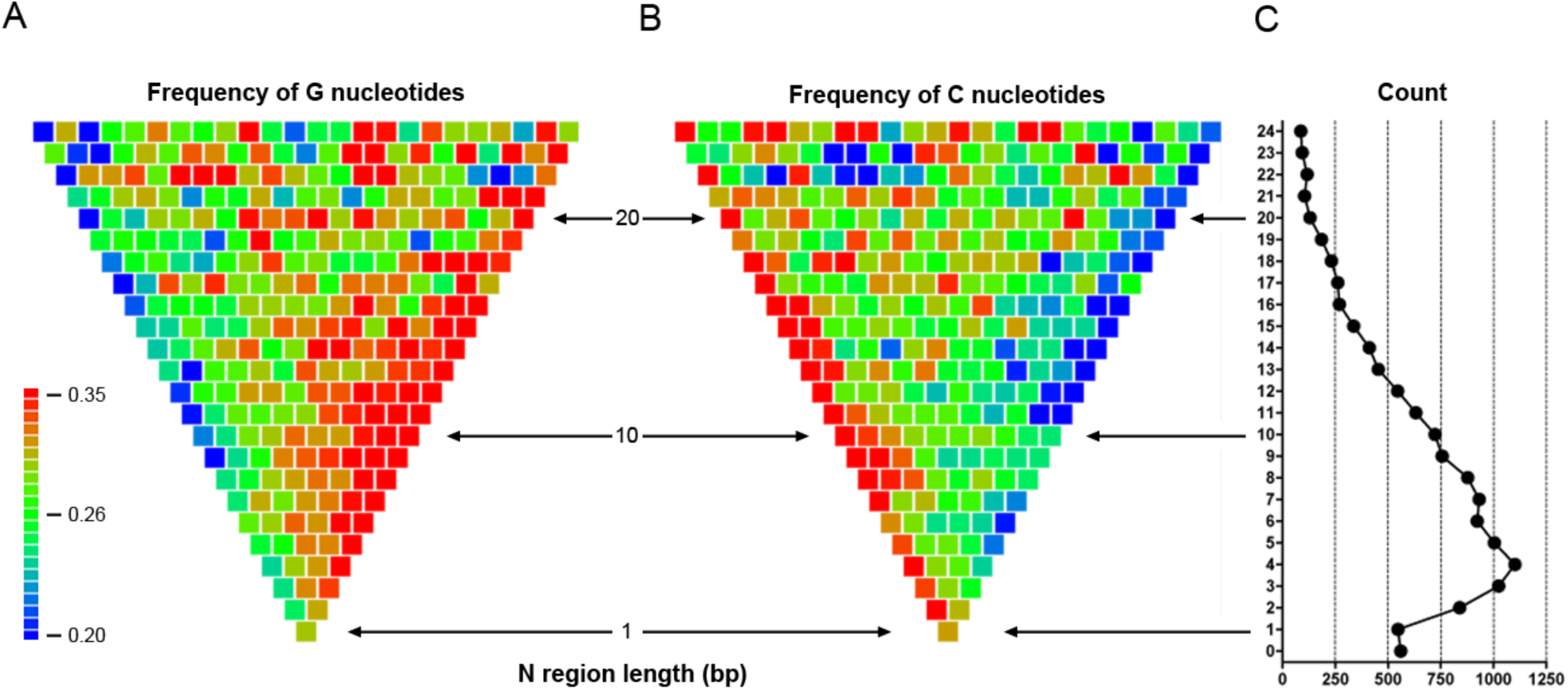
G and C gradients in opposite directions throughout the spectrum of N2 lengths. Only sequences with exactly one identified D gene were included. **A)** and **B)** Heat maps showing the frequencies of the nucleotides G and C, respectively, at specific positions (plus strand) of the N2 region in 3,438 unmutated NP V_H_DJ_H_-rearrangements and 9,061 sterile DJ_H_-rearrangements. Out of these 12,499 sequences, 543 (4.3%) completely lacked N2. Further, 525 (4.2%) with N2 lengths above 24 nucleotides (25-74 nucleotides) were omitted from the figure. Each square represents one base position in the N2 region of a given length from the 5’ to the 3’ end of the region (left to right). The color indicates the frequency of the respective nucleotide. **C)** The corresponding number of sequences of each N region length.

**Figure 2.**
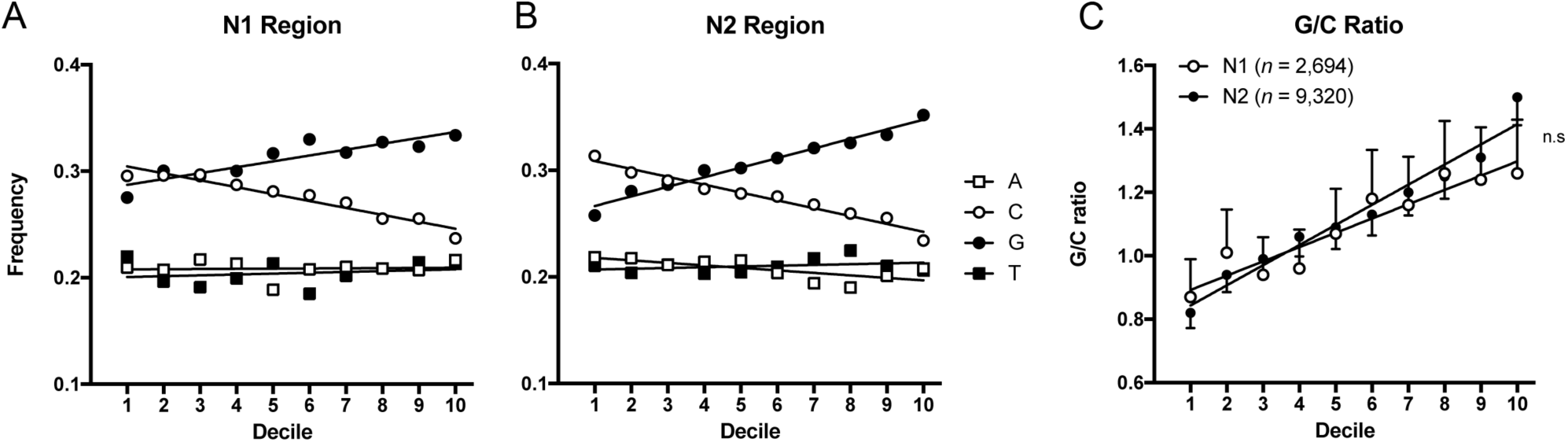
The nucleotide usage across 2,694 N1 regions and 9,320 N2 regions in unmutated NP rearrangements. **A**) A distinct linear decrease was observed for C frequencies across N1 regions (*R*^2^=0.92, *β*=−0.007 [−0.009;−0.004], *p*<0.0001) while a linear increase in G was seen (*R*^2^=0.80, *β*=0.006 [0.003;0.008], *p<*0.0001). **B**) Similar gradients were found in the N2 regions where the C gradient (*R*^2^=0.96, *β*=−0.007 [−0.008;−0.006], *p*<0.0001) was indistinguishable from that in N1 (*p*=0.488), while the G gradient (*R*^2^=0.97, *β*=0.009 [0.008;0.010], *p*<0.0001) differed from that in N1 by a greater slope (*p*=0.005). The dATP gradients differed slightly between N1 and N2 (*p*=0.021). The dATP remained constant throughout N1 (*R*^2^=0.004, *β*=0.0002 [−0.0017;0.0020], *p*=0.863) while it decreased slightly for N2 and (*R*^2^=0.52, *β*=−0.002 [−0.003;−0.001], *p*<0.0001). The dTTP frequencies remained constant throughout both N1 (*R*^2^=0.04, *β*=0.0008 [−0.0010;0.0026], *p*=0.393) and N2 (*R*^2^=0.11, *β*=0.0007 [−0.0003;0.0017], *p*=0.164) and did not differ between N1 and N2 (*p*=0.941). **C**) The G/C ratios across the N regions did not differ between N1 and N2 (*p*=0.054).

### The G/C ratio in N regions was influenced by selection but not somatic hypermutation

The G/C gradients in N1 and N2 depended somewhat on the mutational and functional status of the rearrangements (Figure 3A and B). Functionality clearly had a greater effect on the G/C ratio across N1 regions compared to N2 regions. The increase in G/C ratio was due to a relative enrichment of G in the N1 region (Figure 3C) reaching 34.6% in unmutated P rearrangements compared with unmutated NP rearrangements (30.4%, *p*<0.0001). While significant, the increase in G in unmutated P rearrangements was less pronounced in N2 (Figure 3D). No significant differences in G/C ratio were found for N1 or N2 when comparing mutated P and unmutated P rearrangements (p>0.33).

**Figure 3.**
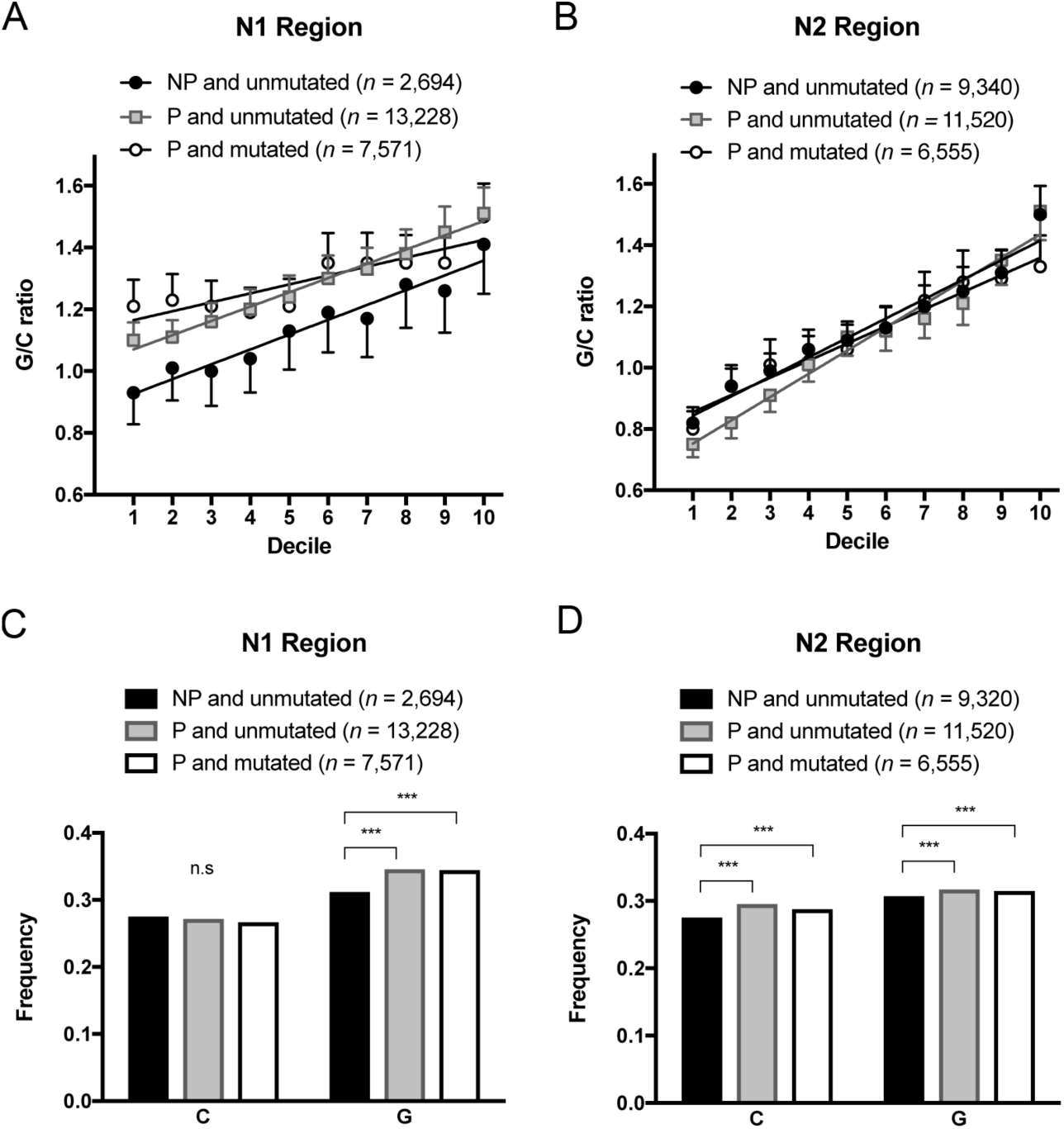
The G/C gradient in N regions in unmutated NP, unmutated P and mutated P rearrangements, respectively. **A**) For N1, the slopes (G/C gradients) did not differ between unmutated NP and unmutated P V_H_DJ_H_-rearrangements (*p*=0.895) but the Y-intercept was lower for unmutated NP (*p*=0.002) **B)** For N2 regions, no differences were found (*p*=0.30). **C)** In N1 the frequency of C nucleotides did not differ significantly between unmutated NP, unmutated P and mutated P rearrangements (*p*=0.084), while the content of G nucleotides was significantly higher in unmutated P and mutated P compared to unmutated NP rearrangements (*p*<0.0001) and **D)** In N2 the frequency of both C and G nucleotides were slightly higher in unmutated P and mutated P compared to unmutated NP rearrangements (*p*<0.0001).

### No significant contribution by secondary rearrangements to the composition of N1

The slight difference in the G gradient observed between N1 and N2 (Figure 2A and B) led us to test whether footprint motifs templated by the 3’-end of a V_H_-gene (putative remains after secondary rearrangements) were present in the sequence material as such footprints may bias the nucleotide distribution in N1. All possible 7-mer, 6-mer, and 5-mer motifs present in the coding part of any of the known germline V_H_-gene alleles between the putative cryptic RSS and the canonical RSS heptamer were identified (Supplementary table 1). For each of them, the number of occurrences was compared between N1 and N2 from all unmutated NP rearrangements with N regions long enough to accommodate the motif. No significant difference was found for any 7-mer or 6-mer motif and only a single 5-mer (CGGAT) reached significance (*p*=0.02, Holm-Bonferroni corrected value). However, this motif was only found in 1.2% of N1 regions and removing these rearrangements did not influence the G, C or G/C gradients in N1 significantly (data not shown). The presence of footprints could therefore not explain the difference observed between the G gradients in N1 and N2.

### Gene segments were subject to different levels of trimming

In rearrangements, the V_H_, D and J_H_-gene segments are often trimmed compared to the germline sequence. Trimming takes place during the joining event of the germline encoded gene segments. In NP V_H_DJ_H_- rearrangements we found that only 68% of V_H_-3’-ends were trimmed, which was significantly less compared to the D-5’-ends (84%), and of D-3’-ends (83%) and J_H_-5’-ends (88%) found in NP V_H_DJ_H_- and sterile DJ_H_-rearrangements (*p*<0.0001, Figure 4). The same pattern was found in P rearrangements. Thus, the N1 region was flanked by an untrimmed gene segment more frequently than N2 (*p*<0.0001).

**Figure 4.**
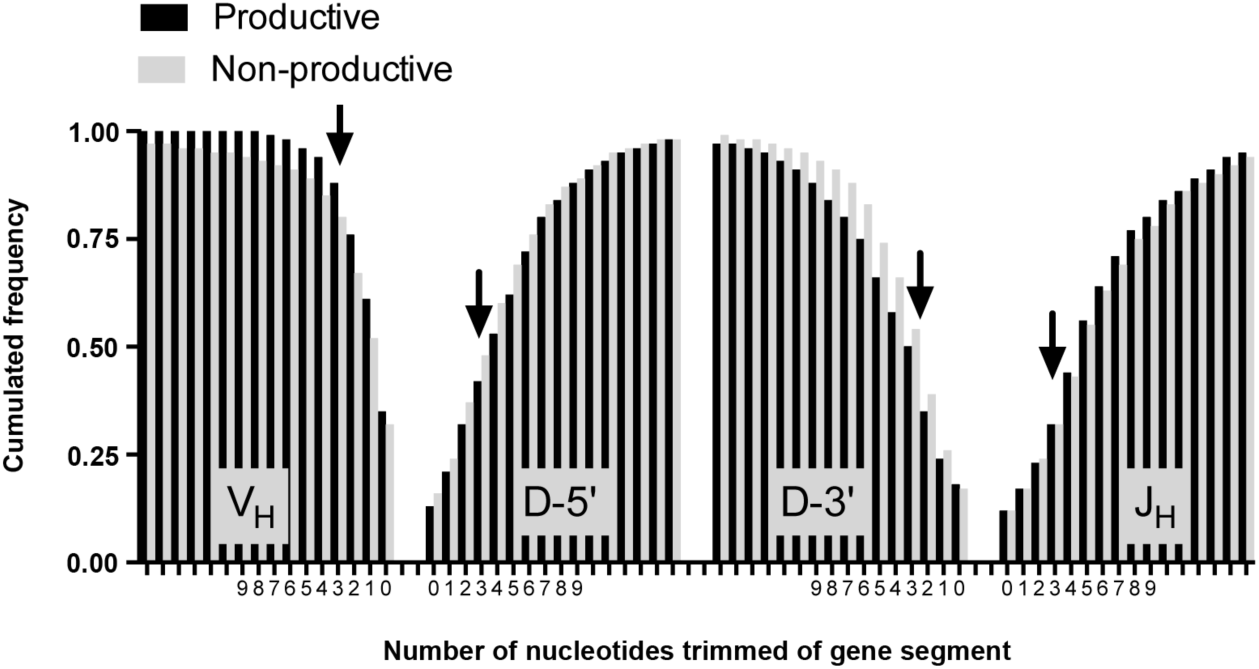
The figure illustrates the number of nucleotides trimmed of gene segments in 29,392 P V_H_DJ_H_-rearrangements and 13,769 NP DJ_H_- and V_H_DJ_H_-rearrangements irrespective of mutation status. The trimming status is depicted as cumulated frequencies of gene segments with 0, 1, 2, 3 or more (up to 15) nucleotides removed from the ends facing N1 (V_H_ and D-5’-end) or N2 (D-3’-end and J_H_). For NP rearrangements, 68% of V_H_-gene segments had been subject to some level of trimming while the same was true for 84% of the 3’-ends of D-gene segments, 83% of the 5’-ends of the D-gene segments and 88% of J_H_-gene segments. Also in P rearrangements fewer V_H_-gene segments (65%) had been subject to trimming compared to the 3’-end (87%) and 5’-end (82%) of D-gene segments and J_H_-gene segments (88%).

### G/C ratios were dependent on gene segment trimming

We next studied the effect of gene segment trimming on the G/C gradient to see whether it was influenced by trimming of the flanking gene segments. This analysis was most informative for the N2 regions where 9,320 unmutated NP V_H_DJ_H_- and DJ_H_-rearrangements had evaluable N2 regions that could be broken down to 6,924 with trimmed gene segments at both ends, 879 with only the D-gene trimmed, 1,352 with J_H_-gene trimming only, and 165 with untrimmed gene segments at both ends. In N2 regions where both flanking gene segments had been trimmed, we found a striking G/C gradient (*R^2^*=0.96, *β*=0.08 [0.068;0.085], *p*<0.0001) (Figure 5b). Interestingly, the G/C ratio changed considerably in proximity of an untrimmed gene segment. In deciles closest to trimmed D-3’-ends, the G/C ratio remained low while it remained high throughout N2 regions if only the J_H_-end was trimmed. This strongly suggested that the G/C gradient observed across the N2 regions was dependent on the trimming of both flanking gene segments. An intriguing observation was inversion of the G/C gradient (*R^2^*=0.65, *β*=−0.07 [−0.12;−0.01], *p*=0.023) seen for N2 regions flanked by two untrimmed gene segments (Figure 5D).

**Figure 5.**
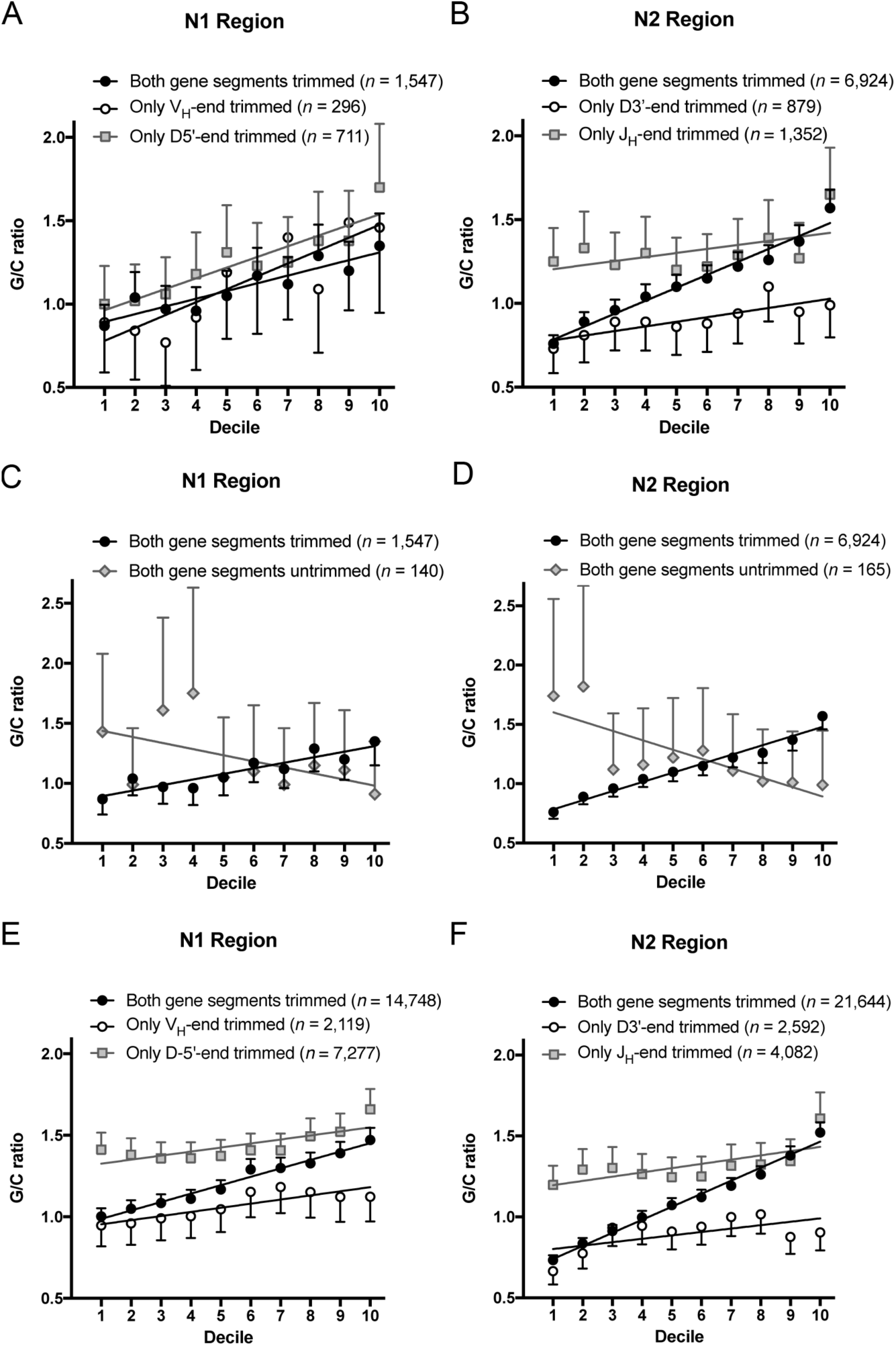
The effect of gene segment trimming on the G/C ratios across N regions in unmutated NP rearrangements (panels A-D) or all rearrangements irrespective of functionality and mutation (panels E-F). **A)** A positive G/C gradient was seen for 1,547 N1 regions flanked by two trimmed gene segments (*R^2^*=0.84, *β*=0.05 [0.03;0.06], *p*<0.0001), the 711 N1 regions flanked by a trimmed D5’-end and an untrimmed V_H_-gene (*R^2^*=0.84, *β*=0.06 [0.03;0.09], *p*<0.0001) and 296 N1 regions flanked by a trimmed V_H_-gene segment and an untrimmed D5’-end (*R^2^*=0.76, *β*=0.07 [0.03;0.11], *p*=0.0004). The slopes did not differ significantly from one another (*p*=0.16) **xB)** A positive G/C gradient was found for 6,924 N2 regions flanked by two trimmed gene segments (*R^2^*=0.96, *β*=0.08 [0.068;0.085], *p*<0.0001) and with a smaller slope for 1,352 N2 regions flanked by an untrimmed D3’-end and a trimmed J_H_-gene segment (*R^2^*=0.66, *β*=0.03 [0.01;0.05], *p*=0.004). No significant G/C gradient was observed in the 879 N2 regions flanked by a trimmed D3’-end and an untrimmed J_H_-gene segment (*R^2^*=0.33, *p*=0.067). The three lines differed significantly from one another (*p*=0.0003) **C)** There was no significant G/C gradient observed in the 140 N1 regions flanked by two untrimmed gene segments (*R^2^*=0.89, *p*=0.15). **D)** There was a negative G/C gradient observed in the 165 N2 regions flanked by two untrimmed gene segments (*R^2^*=0.65, *β*=−0.07[−0.12;−0.01], *p*=0.023). **E and F)** Including all rearrangements regardless of mutation and functionality, we found similar patterns for N1 and N2 both showing significant positive G/C gradients for all three combinations of trimming (N1: *p*<0.002; N2: *p*<0.0001) but with significantly different slopes (N1: *p*<0.0001; N2: *p*<0.0001).

A similar positive G/C gradient was noted for N1 in 1,547 NP unmutated V_H_DJ_H_-rearrangements which were trimmed at both the V_H_-gene and the 5’-end of the D-gene (figure 5A) (*R^2^*=0.84, *β*=0.05 [0.03;0.06], *p*<0.0001). Rather few sequences (*n*= 711 and 296) were available for studying the effect of trimming at one end only, and no significant difference was seen between G/C gradients of sequences trimmed at either end (Figure 5A). However, when all rearrangements, including mutated P rearrangements, were included, similar patterns and significantly different G/C gradients were observed for both N1 and N2 regions dependent on trimming status (figures 5E and F), suggesting that the lack of significance for N1 regions in NP rearrangements could be due to small sample sizes.

We further found that not only were the G/C ratios across N regions influenced by gene segment trimming, they were also influenced by the extent of gene segment trimming (Figure 6). In N2 regions from 11,012 unmutated NP V_H_DJ_H_- and DJ_H_-rearrangements, we found that the high G/C ratio observed in N2 regions trimmed at the J_H_-gene segment and low G/C ratio found in N2 regions trimmed at the D-gene segment (figure 5B) required removal of 3-4 bases from the respective gene segments (Figure 6). In fact, the pattern reversed if the gene segments were trimmed less or remained untrimmed.

**Figure 6.**
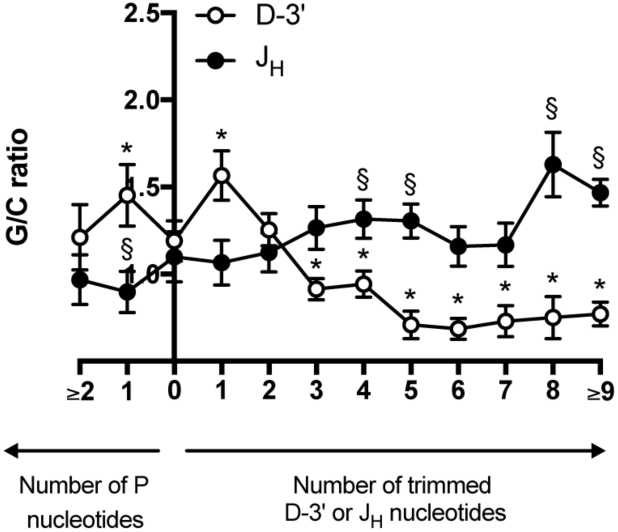
Fine mapping of the impact of gene segment trimming on the G/C gradient across N2 regions. The average G/C ratios in deciles 2-4 and 7-9 were calculated for all rearrangements with N2 segments between 5-38 bp in length and related to number of nucleotides removed from the 3’-end of the germline segment: D-3’ and J_H_, respectively. The elevated G/C ratio observed close to a trimmed J_H_-gene segment required removal of at least 4 bases compared to the G/C ratio when no nucleotides were trimmed. The lower G/C ratio found close to a trimmed D-gene segment required removal of at least 3 bases compared to when no nucleotides were trimmed. * and § indicate where trimming leads to a significant difference in G/C ratio (*p*<0.05) compared to no trimming, for J_H_ and D-3’, respectively.

## Discussion

N addition is an important contributor to junctional diversity and it extensively broadens the human antibody repertoire by usually encoding several amino acids in the central and most important part of the antigen binding site (11). Studies of TdT knockout and TdT transgenic mice have shown that almost all N additions depend on the presence of this enzyme (4, 12, 13). Human TdT shows non-templated polymerase activity *in vitro* and it is exclusively expressed in pro- and pre-T- and B cells, at the site and points in time where N addition occurs (31–33). However, other polymerases capable of non-templated nucleotide addition are present in the cell (e.g. polymerase µ and polymerase λ) and though they are assumed to participate primarily in templated filling of single-stranded segments during NHEJ, it cannot be excluded that one or both contribute to N addition in humans. Further functional studies are needed to define the molecular processes leading to N addition, but only studies of *in vivo* generated rearrangements can elucidate the net result of this complex machinery. Sequencing of *in vivo* generated V_H_DJ_H_-rearrangements have been performed since the 1980s and contributed considerably to our understanding of the generation of diversity. Due to the stochastic nature of N addition, however, informative studies of the fine structure of N addition require large sequence materials only practically available after the introduction of next-generation-sequencing. Moreover, correct interpretation of data requires validated algorithms to define N regions and elimination of templated segments like P nucleotides from the regions. It is also necessary to exclude functional sequences which may have been subjected to selection as well as somatically hypermutated rearrangements in which the base composition may have been changed. Sterile DJ_H_-rearrangements are ideal for these studies because they are non-coding and neither subjected to selection nor somatic hypermutation (27). We analyzed a large dataset composed of DJ_H_-rearrangements yielding unbiased insights into the nature of N2 regions and supplemented with N1 and N2 regions derived from unmutated and non-productive V_H_DJ_H_-rearrangements from several sources which contained only one identifiable D-gene segment. For N1 regions, we also had to consider the possible bias related to templated sequences – so called footprints – which occasionally occurs as a result of a secondary rearrangements (34–38). We have previously shown that such V_H_-gene remnants are too rare to be detectable in the normal B-cell receptor repertoire (26). Here we tested for footprints after secondary rearrangements on this very large sequence material. Again we found that secondary rearrangements are quite rare and unlikely to affect our conclusions in this paper.

### Selection of B cells with functional rearrangements favors G in the N1 regions and less pronounced G and C in N2 regions

Our demonstration that the content of G is higher in N1 regions from productive rearrangements stresses the importance of careful selection of rearrangements for studies of N region synthesis. Moreover, it suggests that some amino acids encoded by G-containing codons (e.g. glycine) are favorable in the CDR3.

### Human N region formation usually occurs by concatenation of two single-strands growing from the 3’-ends of flanking gene segments

The current understanding of the process leading to N additions claims that TdT in a largely stochastic manner adds nucleotides to the 3’-end of the adjacent gene segments with or without prior truncation of the ends and with different preferences for the four nucleotides and a tendency for homo-polymer formation. It is, however, unclear to what extent synthesis occurs on the sense or the antisense strand or whether both strands are involved in the same cell. It is also unknown how truncation of the 3’-ends affects N-region synthesis. The large and highly validated sequence material in this report allows us to discriminate between some of these possibilities. The linear gradients demonstrated in figure 2 cannot be explained by a model assuming N addition only occurring in the plus strand (or the minus strand) by a polymerase with fixed preferences for the individual nucleotides. Neither would a mixture of rearrangements, formed either in the plus strand or in the minus strand, generate such gradients. In both cases, the G/C ratio would be constant over the entire N region. However, before dismissing the single-strand polymerization hypothesis completely, we considered the possibility that homo-polymeric tracts in N regions could be due to nucleotide stacking, as suggested by Gauss and Lieber (39), and that stacking of G’s in the plus strand might generate the observed gradient. *In silico* analyses demonstrated, however, that a gradient of this magnitude could only be generated using extreme levels of stacking (i.e. G following G much more often than any other nucleotide) leading to paucity of short G homo-polymers and preponderance of very long homo-polymers (often as long as the N region) incompatible with our observations (data not shown). In contrast, our data are compatible with a scenario in which two similarly constructed 3’-overhangs, one giving rise to the 5’-end of the N region (plus strand) and the other to the 3’-end (minus strand), generate a gradient balanced around the site where the two 3’-overhangs are joined by microhomology (Figure 7). Assuming a random site for the initial base pairing by microhomology, the concatenation hypothesis would predict the linear gradients that we found. Alt and Baltimore (11) suggested that N regions arise by concatenation of two non-templated strands growing from the 3’-ends of the rearranged gene segments. In 1996, Kepler and colleagues (24), however, studied the nucleotide composition of N1 regions in 543 murine NP V_H_DJ_H_-rearrangements statistically and found the base composition incompatible with the concatenation hypothesis. Based on a calculated homogeneity index they suggested that N regions are primarily formed by a single-strand growing from one of the gene segments (i.e. the single-strand polymerization hypothesis). The discrepancy between our conclusions and that of Kepler *et al.* (24) could be due to differences between mice and humans. However, several issues are problematic with using the homogeneity index to disqualify the concatenation hypothesis for murine N regions. The index measures the tendency of a homogenous pair of the nucleotides G or C (G..G or C..C) in any two positions in the N region separated by n-1 nucleotides compared with any possible pairing (G..G, G..C, C..G, or C..C). An index below 1/2 favors concatenation, when n is large, because the two nucleotides will tend to represent bases added on opposite strands. Kepler *et al.* found values above 1/2 in their N region material (24). The method is, however, based on an assumption of statistical independence of nucleotides added by TdT, but several studies suggest an inherent property of TdT to generate homopolymeric tracts due to base stacking (39). With the short nature of murine N regions, ranging from 1-7 nucleotides in length (3, 12, 40) the sensitivity of the method is low and stacking alone may explain high values for the homogeneity index for closely positioned bases. Moreover, the sensitivity of the homogeneity index is not known because it depends on the ratio between the frequencies with which G and C are added during polymerization *in vivo*, which is currently unknown. When we calculated the homogeneity index for our N2 sequences, we did not find high values (above 1/2) for all values of n as predicted by the single-strand polymerization hypothesis. Rather values below 1/2 were often seen for large values of n (Supplementary figure 2). We conclude that our data strongly suggest that human N1 and N2 regions are usually formed by concatenation of two opposite strands growing from the 3’-ends of the adjacent gene segments.

**Figure 7.**
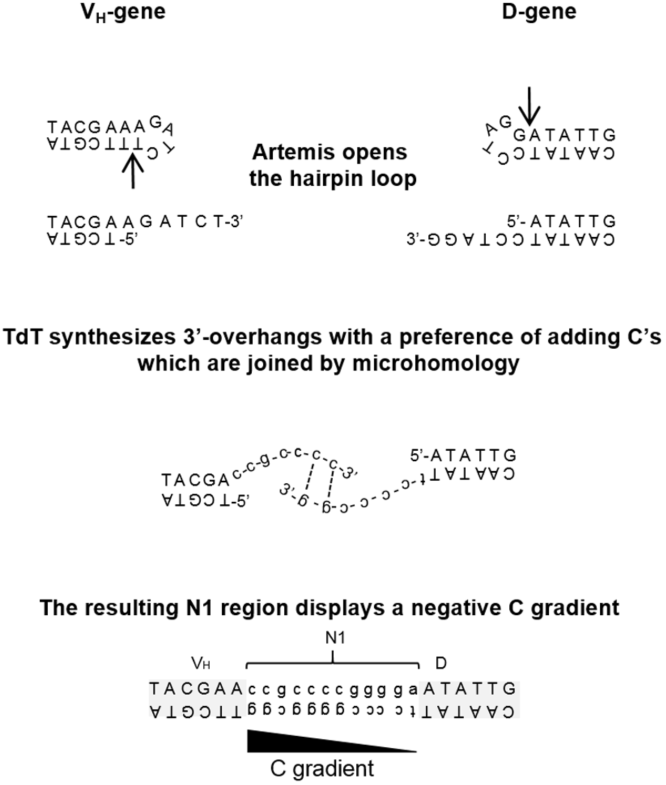
Graphic sketch illustrating the process by which the C gradient is generated during joining of the V_H_-gene and D-gene segments. First Artemis nicks the hairpin loop generated by the RAG-complex resulting in a 3’-overhang. This 3’-overhang is processed by TdT, possibly removing it entirely as depicted in the figure and synthesising an N-addition preferentially composed of C. The two 3’-overhangs are then joined by microhomology and the resulting N1 region will display a negative C-gradient and positive G-gradient.

### The nucleotide most often added during polymerization is C

When accepting the concatenation hypothesis for the majority of rearrangements, it follows inevitably from our data that the nucleotide most often added during polymerization is C followed by G whereas A and T are added at similar, lower frequencies (A slightly more often than T). Assuming that the bases are added by TdT, our data indicate that human TdT preferentially adds Cs during N addition rather than G *in vivo*. This is clearly at odds with the current understanding. Nevertheless, Kepler and colleagues (24) also observed a trend for higher G content in the 3’ end of 543 murine N1 regions and Collins *et al.* more recently noticed a similar G/C gradient in N2 (but not N1) in 199 human V_H_DJ_H_ sequences (41). Both considered this an argument against concatenation, however, because of the preference of TdT for G as demonstrated *in vitro* which, if replicated *in vivo*, would predict the opposite result. It is true that early *in vitro* studies have demonstrated that purified TdT from calf thymus adds G at a significantly higher frequency (60-70%), than C (9-17%), T (11-16%) and A (2-9%) when dNTPs are provided in equimolar concentrations (15). However, *in vivo* dNTPs are not present in equimolar concentrations and the cellular levels of individual dNTPs in different cellular compartments could therefore be responsible for the differences in dNTP usage during the N region formation (42, 43). Notably G concentrations have been shown to be considerably lower than C concentrations in the nucleus of HeLa cells (43). Furthermore, studies in human thymocytes have shown that even though the levels of G increase during the S phase of the cell cycle, they are three fold lower than the C levels during the G_0_/G_1_ phase in which V(D)J rearrangement occurs (44). The tendency to add nucleotides may also depend on other proteins which the TdT enzyme interacts with *in vivo*. For example, human DNA-PKcs has been found to dramatically change bovine TdT’s preferences for individual nucleoside triphosphates (23). Based on our data, we therefore suggest that during *in vivo* rearrangement in human pro- and pre-B cells, C is the most frequently used substrate by TdT followed by G, A, and T in that order.

### Gene segment trimming affects the G and C content in final N region

A novel finding of this study is that single strands growing from trimmed gene segments tend to contribute more to the final N region when the other end is untrimmed. This implies that the N region has been synthetized primarily in one direction in these situations. A possible explanation is that the short isoform of TdT with polymerase activity, in complex with other proteins at the coding end, works more efficiently at trimmed ends *per se* or that its recruitment to the complex is dependent on prior recruitment of factors with exo- (long isoform of TdT) or endonuclease (Arthemis) activity. If this is the case, TdT activity may sometimes be delayed at one gene segment and *de facto* be unidirectional (Figure 7). Our finding that the tendency for unidirectional synthesis is most extreme when the gene segment is trimmed ≥3 nucleotides from the heptamer could suggest that the coding end is blunt-ended before TdT transferase activity is initiated, because Artemis often nicks 2-3 bases from the hairpin loop. However, the existence of V_H_DJ_H_-rearrangements with two untrimmed gene segments argues against strict dependency of TdT’s transferase activity *in vivo* on prior nuclease activity. Another possibility is that concatenation of two growing single-strands occasionally finds microhomology in unprocessed 3’-overhangs generated by Arthemis which would also give rise to the observed patterns. Our data do not allow us to discriminate these two scenarios.

Our data forces us to revise the current understanding of TdT’s preference for G during N addition *in vivo*. We suggest the explanation is to be found in the complex machinery operating on the trimmed gene segment ends during rearrangement, which may differ from the conditions present *in vitro*. The fact that nucleotide preferences of TdT may depend on the situation is evident from the data of Gangi-Peterson *et al*. who found significantly different G and C contents in N regions occurring in coding joints compared to signal joints (25). In fact, the tendency to add G at the 3’-end of signal joints was not found in the coding joints where a trend for preponderance of C was actually observed. Signal joints differ from coding joints by lack of hair pin loop formation and trimming, while coding joints are usually trimmed. This is reminiscent of our finding in this paper of positive G/C gradients in N regions between trimmed gene segments, but negative G/C gradients in the relatively few rearrangements with two untrimmed gene segments. Thus, it is possible that TdT changes its nucleotide preferences dependent on whether the gene segment is trimmed or not, most likely due to differences in the protein complexes formed. This may not only explain our findings, but also explain differences between N region composition of signal joints versus coding joints. More research is needed before we fully understand the intricate mechanism of N addition, but focus should be on mechanisms that can explain the fine structure of N additions revealed in this paper.

